# Synchronised oscillations in growing cell populations are explained by demographic noise

**DOI:** 10.1101/2020.03.13.987032

**Authors:** Enrico Gavagnin, Sean T. Vittadello, Gency Gunasingh, Nikolas K. Haass, Matthew J. Simpson, Tim Rogers, Christian A. Yates

## Abstract

Understanding synchrony in growing populations is important for applications as diverse as epidemiology and cancer treatment. Recent experiments employing fluorescent reporters in melanoma cell lines have uncovered growing subpopulations exhibiting sustained oscillations, with nearby cells appearing to synchronise their cycles. In this study we demonstrate that the behaviour observed is consistent with long-lasting transient phenomenon initiated, and amplified by the finite-sample effects and demographic noise. We present a novel mathematical analysis of a multi-stage model of cell growth which accurately reproduces the synchronised oscillations. As part of the analysis, we elucidate the transient and asymptotic phases of the dynamics and derive an analytical formula to quantify the effect of demographic noise in the appearance of the oscillations. The implications of these findings are broad, such as providing insight into experimental protocols that are used to study the growth of asynchronous populations and, in particular, those investigations relating to anti-cancer drug discovery.

**Statement of Significance:** Recent experiments have reported strong evidence of periodic oscillations in the proportion of young and old melanoma cells. The biological mechanism generating this synchronisation and the potential impact that can have on commonly used experimental protocols is still unclear. Here we studied a population of melanoma cells for which we found oscillations in the proportions of cells in each phase of the cell cycle. We demonstrate that these observations may be triggered by intrinsic demographic noise alone, rather than any active synchronisation mechanism requiring cell-cell communication. Our findings may have implications for typical experimental protocols which aim to produce asynchronous cell populations.

## I. INTRODUCTION

Growing populations are a crucial feature of many biological phenomena, from the clonal expansion of cancer cell lines to the increase in numbers of infected individuals during a disease outbreak. A deeper understanding of cell proliferation sheds light on a vast range of biological processes, from morphogenesis to tumour growth [1, 2], understanding and predicting the time evolution of these growing population is, therefore, of fundamental biological relevance [3, 4]. The initial stages of growth in both these scenarios are typically considered to be exponential as cells replicate without restriction or disease spreads into an otherwise susceptible population.

Standard mathematical modelling approaches assume that cell divisions are independent events with exponentially distributed waiting times. This gives rise to exponential growth in unstructured populations [5]. In cell biology, this approach has been supported by classic experimental studies for large populations under favourable growth conditions [6, 7]. However, when smaller populations are considered - for example clones of a single progenitor cell - the classical model of exponential growth fails to capture the variable per capita growth rates caused by non-exponentially distributed cell cycle times and more sophisticated models are necessary [8–12].

Due to recent technological advances, we are now able to access accurate data revealing the structure of dynamic cell populations [13, 14] using, amongst other tools, proliferation assays: an *in vitro* experimental protocol in which the growth of cell populations is monitored over time [15]. In particular, in recent work [16], we assayed the proliferation of melanoma cells labelled with FUCCI (Fluorescent Ubiquitination-based Cell Cycle Indicator [17, 18] - see Fig. 1 and Section II B of Materials and Methods) which allowed us to track the number of cells in particular phases of the cell cycle over a timespan of 48 h. Strikingly, the proportion of cells in the first phase of the cell-cycle, gap 1 (G1), demonstrates clear and unexplained fluctuations during the entire duration of the experiment. This synchrony, reproducible over multiple cell lines and experimental replicates, has potentially serious implications for studies of rapidly dividing cell populations and demonstrates that classical, and widely adopted exponential models of population growth are insufficient to capture either individual-level or population-scale growth dynamics.

**Fig. 1.**
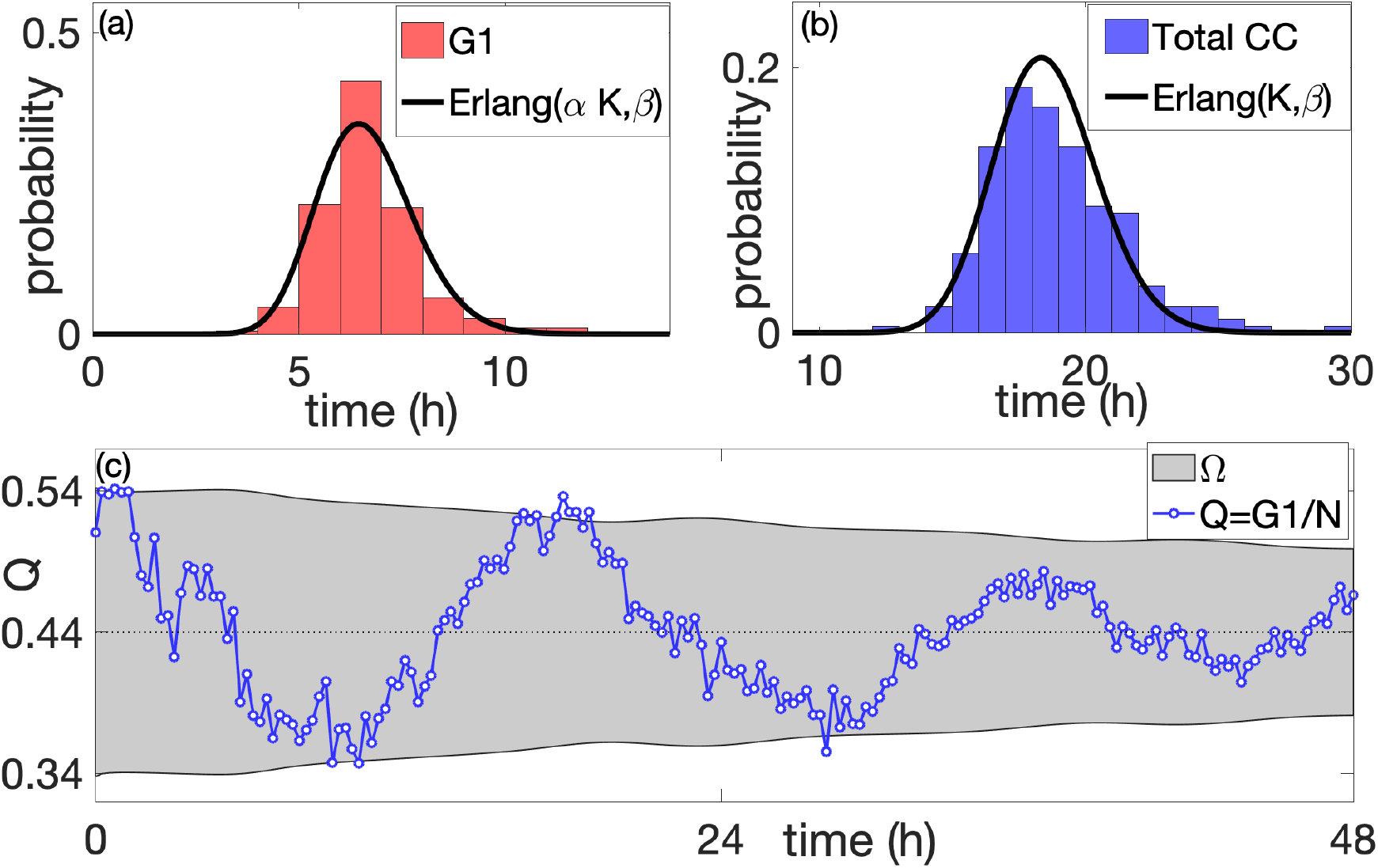
Cell cycle and phase time distributions and synchronised oscillations. Panel (a) shows a comparison between the distribution of the duration of the G1 phase obtained by tracking 200 randomly chosen cells (red histogram) and the Erlang distribution with the same mean and variance (black curve). Panel (b) shows the comparison between the distribution of the full cell cycle time of the same tracked cells (blue histogram) together with the corresponding Erlang distribution. Panel (c) shows one time series trajectory of the proportion of G1 cells, *Q*, obtained from the experiments (blue line), together with the envelope of two standard deviations from the mean (light grey region) predicted using the multi-stage model. The parameters of the multi-stage models are obtained by fitting the distribution of the total cell-cycle time and G1 duration (see Section S.4 of Supplementary Materials): *K* = 92, *αK* = 33, *β* = 4.96*h*^−1^, *v* = 94.3 and *N*_0_ = 381

Inspired by our experimental findings, we build a multi-stage mathematical model for cell proliferation which represents the cell-cycle as a series of discrete stages [16]. The waiting time distribution between consecutive stages is exponential, meaning that the cell cycle time (CCT) follows a more general class of distributions, known as hypoexponential Yates *et al*. [8]. This family of distributions has been shown to provide good agreement with the experimental cell-cycle time distribution data [8, 13, 14, 19]. By deriving a deterministic representation of the population dynamics under the multistage approach we reproduce the cell-cycle fluctuations observed in the experiments. This suggests that multi-stage models are a suitable framework for investigating the phenomenon of cell-cycle synchronisation. However, since in [16] the parametrisation of the model is carried out individually for each experimental trajectory, our previous study did not explain the origin of such oscillatory phenomena nor their asymptotic behaviour.

In a growing cell population, cell division acts as a natural source of synchronicity by increasing the number of phase-synchronised cells. This progeny form of synchronicity plays a fundamental role when the population size is small (for example in the case of a single-cell lineages). As the population grows, the effect of initially synchronous progeny gradually decreases and it eventually becomes negligible for a large population, at which point the phase distribution - the distribution of cells in each phase of the cell cycle - reaches an invariant state and the total population grows exponentially (*asymptotic* regime). When a small population is sampled from a larger one, for example during a typical cell proliferation assay, the intrinsic randomness of the process plays a dual role. Stochastic finite-size effects can lead to the sampled population being far from the invariant distribution. That sampled population will have a synchronised oscillatory state. However the individual variations in the cell-cycle time tend to break the correlation of synchronised cells, resulting in a gradual desynchronisation the population (*transient* regime). The presence of these two regimes (a transient-oscillatory regime and an asymptotic-exponential regime) is a common feature of many structured growing population [9, 10, 20]. It is not surprising, therefore, that these two phases play distinct but critically important roles in the dynamics of a growing cell population.

The extent to which this natural transient phase is responsible for the observed oscillatory phenomenon remains unclear. Our goal is to design a suitable theoretical framework, based on the multistage model (Section II D), which allows us to quantify for a given population the magnitude of synchronicity triggered and sustained by demographic noise - a general term which describes to the randomness emerging at the level of a finite-sized population when individual birth and death events occur stochastically. The focus of our analysis is on quantifying the expected amplitude of the oscillations, as this quantity can be easily measured and directly linked to experimental observations, even when the data are limited to short time windows.

In this study, we establish that strong oscillations observed in growing populations of cells can be triggered by demographic noise alone. This finding demonstrates that it may not be necessary to appeal to an external synchronisation mechanism requiring cell-cell communication to explain synchronisation observed in experiments. To do this we first analyse the multi-stage model with a particular focus on characterising the transient and asymptotic phases. By deriving a stochastic mesoscopic model, we study the effect of stochasticity in the system and obtain an analytical formula that can be used to quantify the amplitude of the fluctuations due to finite-sample effects. Finally, we parametrise the multi-stage model by fitting the G1 and total cell-cycle time distributions, obtained from single-cell tracking data, and compare our predictions with the time series obtained in the experiments.

Our central finding is that the fluctuations in the subpopulation of G1 cells in the proliferation assay are of the same magnitude as those induced by demographic noise alone, which suggests finite-sample effects provide a straightforward, yet often overlooked explanation of the observed synchronisation. Our study examines the specific impact of demographic noise on the dynamics of the amplitude of the oscillations, predicting that the observed synchronicity is a transient phenomenon for which we can predict the corresponding characteristic decay time.

The fact that the observed synchrony is a generated by demographic noise, and not a feature peculiar to the cell line we studied, means that we expect the same phenomenon to be observed in a wide range of other populations undergoing stochastic growth, as exemplified in a number of studies [4, 15, 21–23]. Moreover, the generality of the mathematical models adopted throughout, allows this study serve as a showcase of the potential applications of multi-stage models. The possible implications of our study therefore range from revised experimental protocols, to altered cancer treatment schedules and from new ways of understanding early infection progression within the body to strategies for prevention of the spread dissemination of disease in the early stages of an outbreak. The characterisation of the transient nature of synchronised behaviour may lead the way to new experimental designs for a broad range of experimental protocols in which cell cycle synchronisation is of vital importance [24].

## II. MATERIALS AND METHODS

### A. Cells and cell culture

The human melanoma cell line C8161 (kindly provided by Mary Hendrix, Chicago, IL, USA) was genotypically characterised [25, 26], grown as described by Spoerri *et al*. [27] and authenticated by STR fingerprinting (QIMR Berghofer Medical Research Institute, Herston, Australia)

We maintain the cell cultures to prevent any induced synchronisation from cell cycle arrest in G1 phase. In general, such induced synchronisation can occur through various experimental conditions, namely contact inhibition of proliferation at relatively high population densities [28], decreased pH of the growth medium due to the concentration of acidic cell-metabolites such as lactic acid [29], and reduced availability of nutrients such as serum [21]. We prevent induced synchronisation by passaging the cells every three days, and on the day prior to setting up an experiment, to maintain a subconfluent cell density and a fresh growth medium, so that the cell culture conditions are never such that they cause G1 arrest.

We note that there are other factors that can induce cell synchronisation. For example, during the suspension, prior to seeding, some cells may die due to detachment - in particular those close to mitosis. Clearly, this phenomenon might lead to further deviation from the invariant distribution and hence, to amplification of the appearance of the oscillations. Since the aim of our study is to quantify the oscillations arising only from finite-sample effects, we do not account for this phenomenon in our model.

### B. Fluorescent ubiquitination-based cell cycle indicator

To generate stable melanoma cell lines expressing the FUCCI constructs, mKO2-hCdt1 (30–120) and mAG-hGem (1–110) [17] were subcloned into a replication-defective, self-inactivating lentiviral expression vector system as previously described [30]. The lentivirus was produced by cotransfection of human embryonic kidney 293T cells with four plasmids, including a packaging defective helper construct (pMDLg/ pRRE), a Rev plasmid (pRSV-Rev), a plasmid coding for a heterologous (pCMV-VSV-G) envelope protein, and the vector construct harboring the FUCCI constructs, mKO2-hCdt1 (30–120) and mAG-hGem (1–110). High-titer viral solutions for mKO2-hCdt1 (30/120) and mAG-hGem (1/110) were prepared and used for co-transduction into eight biologically and genetically well-characterized melanoma cell lines (see above), and subclones were generated by single-cell sorting [18, 27, 31].

### C. Image processing and analysis

The microscopy data consist of multi-channel time-series stacks which are processed and analysed automatically with Fiji/ImageJ and MATLAB as described in Vittadello *et al*. [16].

To obtain the time distribution of the G1 phase cells and of the cell cycle we selected 200 cells towards the beginning of the experiment. To do this, we first labelled all the automatically detected cells on the first frame of the processed merged image of the red and green channels (using the routine Analyse particle of Fiji/ImageJ), we then selected 100 labelled mother cells uniformly, without replacement. For each selected mother cell we manually recorded the time intervals corresponding to the G1 phase (*i*.*e*. between the mitosis event of the mother cell and the first appearance of the cell in the green channel) and to the cell cycle (*i*.*e*. between the mitosis event of the mother cell and the last appearance of the cell in the green channel) of its two daughter cells. We ignored cells which did not reach mitosis before the end of the experiment or move out of the microscopy window (1% of the selected cells).

### D. Multistage agent-based model

We adopt an agent-based model (ABM) for the growth and division of cells, following [8, 13, 19]. In this formulation the cell-cycle is represented as a series of *K* stages through which a cell progresses before it divides. We choose the waiting time to progress from one stage to the next to be exponentially distributed with rate *β*, independent from all other events. When a cell passes through the final stage, it divides into two new daughter cells, both initialised at stage one. This is a simplified model of the cell cycle, however, it is sufficient for the purposes of this study and (as we will show later) it gives a good fit to experimentally observed distributions of cell cycle time.

The *K* stages of our model are grouped into sections corresponding to the known phases of the cell cycle. In particular, we say that a cell is in the G1 phase if it is in one of the first *αK* stages, where *α* ∈ [1*/K*, 2*/K*, …, 1] is a constant to be determined by comparison with data. Expressed as a sum of exponential random variables, the duration of both the G1 and the entire cell cycle are Erlang distributed with parameters (*K, β*) and (*αK, β*), respectively. Fig. 1 (a) and (b) show the maximum likelihood fit of the model simultaneously to both the duration of the G1 phase and the total cell-cycle time for the melanoma cell line C8161. In this example we find parameters *K* = 92, *αK* = 33, *β* = 4.96h^−1^. The measured cell cycle time has an average of 18.5 hours with standard deviation around 2 hours (see Section S.4 of Supplementary Materials).

We define the state vector **X**(*t*) = (*X*_1_(*t*), *X*_2_(*t*), …, *X*_K_(*t*)), where *X*_k_(*t*) denotes the number of cells in stage *k* at time *t*. Our model can be represented as a series of chemical reactions, namely

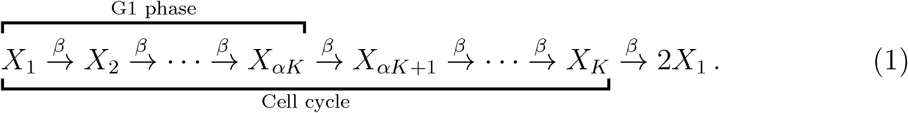

Note that one can write down a system of master equations for the set of chemical reactions above, however this system is analytically solvable only for the simple case of *K* = 1 [8]. We write 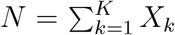 for the total number of cells,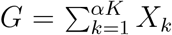 for the number of cells in G1 phase. As the population grows, the proportion of cells in each stage will eventually converge to a fixed value lim_t→∞_ *X*_i_*/N* = *u*_i_, for *i* = 1, …, *N*. Together, these proportions are known as the *invariant stage distribution*, **u**. In Section III B we prove this and derive an exact expression for the limit **u**. On shorter time horizons, the system exhibits transitory oscillations about the invariant distribution [32, 33].

## III. RESULTS

### A. Multistage model recapitulates experimental observations

To assess the amplitude of oscillations, in what follows we develop a mathematical theory for the behaviour of the proportion, *Q* = *G/N*, of G1-phase cells (see Material and Methods). The first part of our analysis reveals long-lived damped oscillations in the expected value of *Q* in a growing population, while the second shows how this effect is initiated and sustained by demographic noise.

Our experimental data are 30 time series of images taken from proliferation assays, as previously reported in Vittadello *et al*. [16] - see Section II A of Materials and Methods and Fig. S.3 of the Supplementary Materials for three snapshots of the microscopy images. Each time series captures a 48 h time window following an incubation period of 24 h. In Fig. 1 (c) we report an example of experimental trajectory of *Q* (blue line) with marked oscillations and the envelope of two standard deviations about the mean, Ω (light grey region) obtained from the multi-stage model. The trajectory shows clear oscillations about the mean (about three complete cycles in the 48 h window) and 94% of the data points lie inside Ω (see Section S.6 of the Supplementary Materials for all the 30 time series of the experiments). The period of the oscillations is approximately one CCT, *i*.*e*. 18.5 h, which is confirmed by the power spectrum analysis (reported in Figure S.5 of the Supplementary Materials). In Fig. 2 (a) we compare the amplitude of the oscillations: we present the time series of the sample standard deviation of *Q* (the proportion of G1-phase cells), *σ*_Q_(*t*), obtained from the 30 experimental trajectories (blue line), together with the corresponding theoretical value (grey line) predicted from the multi-stage model (see Section S.3 of Supplementary Materials). The plot shows that, despite the fluctuation in the experimental data, there is a good agreement between the two time series for the entire duration of the experiment.

**Fig. 2.**
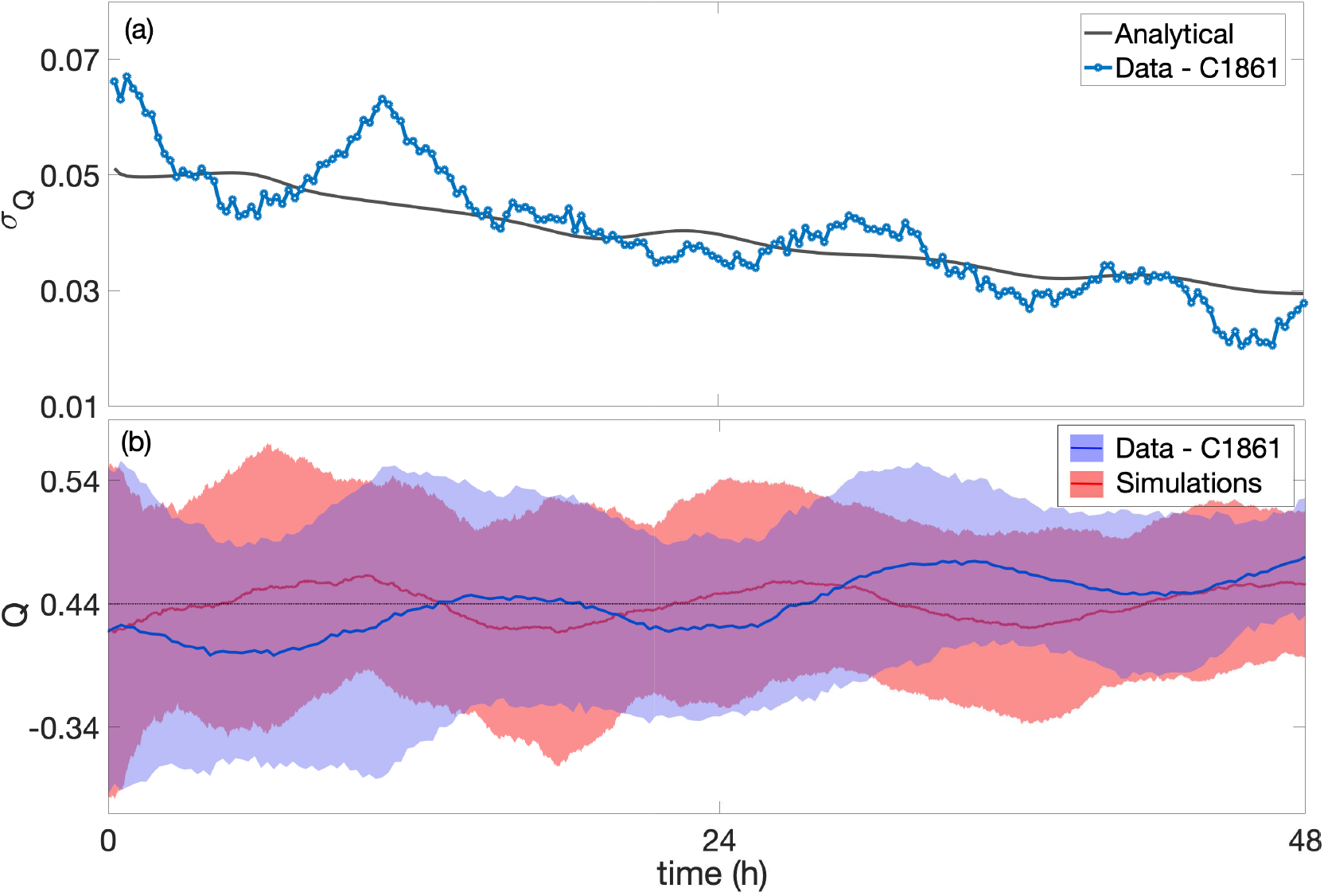
Comparison amplitude of oscillations between experimental data and model prediction. Panel (a): time series of the standard deviation of *Q, σ*_Q_, obtained from the experiments (blue line) and the analytical prediction of the model (grey line). Panel (b): overlay of envelopes of two sample standard deviations (shaded regions) around the sample means (full lines). The blue envelope is obtained from the 30 experimental time series. The red envelope is obtained from 30 independent simulated trajectories. The initial state of the simulations is chosen to match the mean and sample standard deviation of the data. The parameters of the model are the same as in Fig. 1.

In Fig. 2 (b) we display the envelopes of two sample standard deviations around the sample mean of *Q*(*t*), obtained from the 30 experimental time series (blue) and from 30 independent simulations of the multi-stage model (Langevin model - see Section III C) initialised to match the experimental sample mean and standard deviation at time 0. The plot highlights some qualitative similarities between the two envelopes, in particular the rate of decay of the sample standard deviation obtained from simulations is consistent with the experiments. Interestingly, a periodicity of approximately one CCT is visible in the time series of the sample mean which is qualitatively captured in the trajectory obtained from independent simulations, suggesting that this phenomenon may be due to the relative small size of the sample (30 trajectories).

### B. Understanding the transient and asymptotic dynamics

In order to understand the interplay between the transient oscillatory dynamics and asymptotic exponential growth, we begin by writing down the equations governing the dynamics of the expected number of cells in each stage 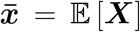. Here *expected* should be interpreted as the average over many experiments with precisely the same initial condition - we will later see that the variability of the initial condition is a different feature that can also lead to the emergence of oscillations. From the model formulation we directly obtain

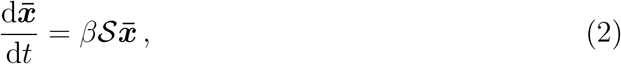

where *β* is the rate of progression through the model stages, and ***S*** is the corresponding stoichiometry matrix. This matrix has non-zero entries ***S***_k,k_ = −1, ***S***_k,k+1_ = 1 for *k* = 1, …, *K* − 1 describing progression between stages, and ***S***_K,K_ = −1, ***S***_1,K_ = 2 describing cell division.

For the purpose of the analysis, we assume *β* = *K* throughout, so that the average cell-cycle time is normalised to unity. The characteristic polynomial of the matrix ***S*** is 𝒫(*y*) = (*y*+1)^K^ −2, from which the eigenvalues of ***S*** are 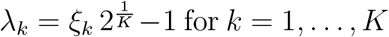 for *k* = 1, …, *K*, where *ξ*_k_ = *e*^2πik/K^ is a *K*-th root of unity. By solving a series of recursive equations, one can write down the left- and right-eigenvectors associated with the *k*-th eigenvalue of ***S***, which we denote **u**^k^ and **v**^k^, respectively. Specifically, we have

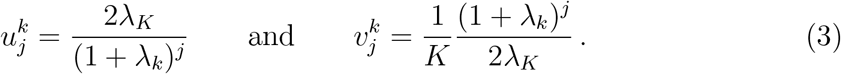

We drop the index *k* whenever we refer to the eigenvalue with maximum real part and the corresponding eigenvectors, i.e. 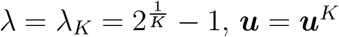, **u** = **u**^*K*^ and **v** = **v**^*K*^.

Notice that from the system of Eq. 2 we can write 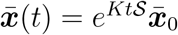, where 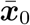 denotes the initial number of cells per stage. In order to study the matrix exponential *e*^KtS^, we first notice that we can write down the (*i, j*) element in terms of the eigenvalues and eigenvectors of ***S*** as

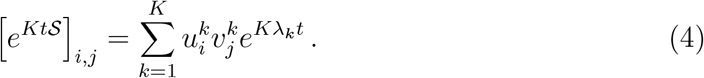

Notice that as *t* → ∞ the leading term of Eq. 4 is *u*_i_*v*_j_*e*^*Kλt*^ and, hence 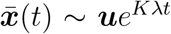 determines the long-time behaviour the system of Eq. 2. We can use this fact to study the limiting behaviour of *Q*: we write 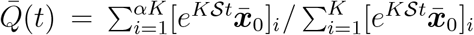 and by looking at the first two terms of 4, we obtain lim_t→∞_ 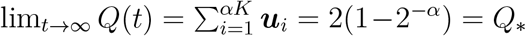.Notice that convergence to *Q*_∗_ occurs with an exponential decay rate given by the spectral gap of the stoichiometry matrix, ℜ[*λ*_*K*−1_] − *λ*_*K*_ (see Fig. 3).

**Fig. 3.**
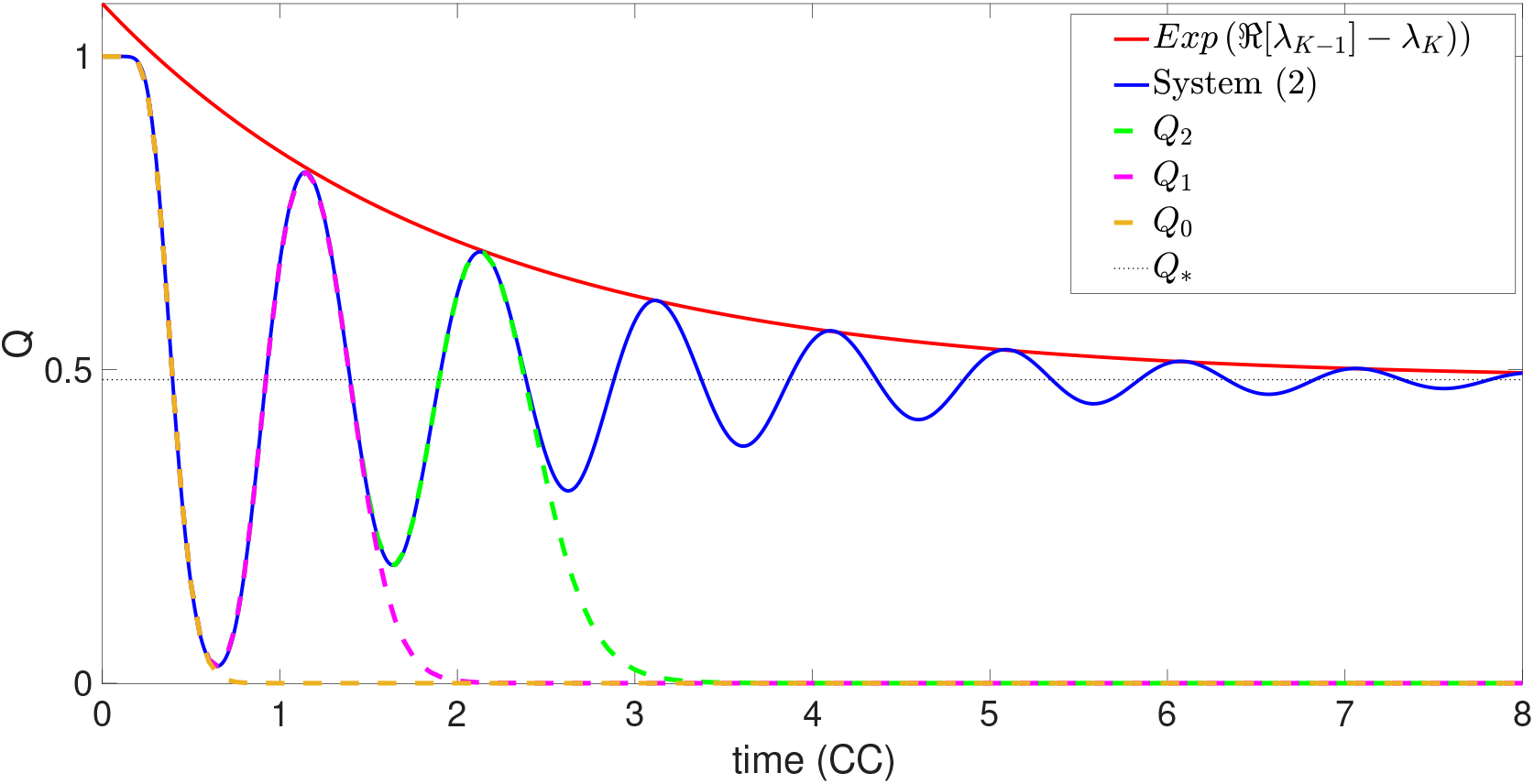
The transient oscillatory dynamics. The figure shows the plot of the ratio *Q*(*t*) obtained by solving the deterministic system of Eq. 2 numerically (blue solid) initialised with **X**(0) = *N*_0_**e**_1_ and parameters *K* = 40, *N*_0_ = 100 and *α* = 0.4. The dashed lines represent the short-time approximation obtained by truncating expression of Eq. 7 up to 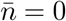 (dashed yellow), 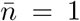 (dashed pink) and 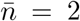 (dashed green). The red solid line shows the exponential decay of the oscillations.

We now focus on the transient behaviour of the system of Eq. 2. We substitute the expressions 3 into the formula 4 and, by exploiting a remarkable identity of the Mittag– Leffler function [34], we transform the finite sum over eigenvalues on the right-hand side of 4 into an infinite sum over the cycles of the oscillatory solutions. Precisely, we write

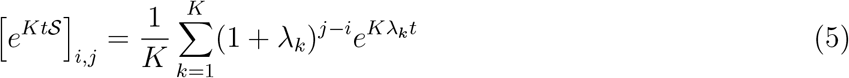

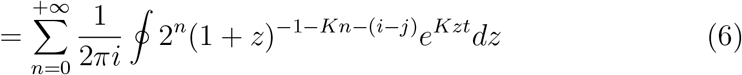

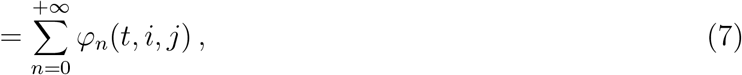

where *ϕ*_n_(*t, i, j*) = 2^n^*e*^−*Kt*^(*Kt*)^*Kn*+i−j^*/*(*Kn* + *i* − *j*)!.

We can now use the expression 7 to approximate *e*^*KtS*^ for short times, by truncating the sum over *n* to a finite index,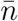. For example, let us consider an initial population of *N*_0_ cells perfectly synchronised at the beginning of the cell cycle, *i*.*e*.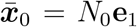.Then we define 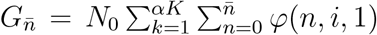 and 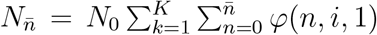. In Fig. 3 we plot 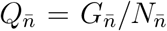, for 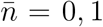and 3, together with *Q* obtained by solving system of Eq. 2 numerically. The plot illustrates how each term of the sum in Eq. 7 contributes one additional oscillation to the transient dynamics of the proportion of G1-phase cells. We now have a complete picture of how oscillations propagate on average in the growing population. It remains for us to show how those oscillations are created and sustained.

### C. Finite-sample effects trigger and amplify oscillations

There are two sources of randomness that are relevant to our model of cell population growth: the finite-size effects involved in the initial sampling, and the individual cell variation of CCT. In order to take into account the stochasticity in the initial population of cells we sample the initial stage distribution, ***x***(0), by drawing from the invariant distribution ***u***. Precisely, the proportion of cells on stage *i* at time 0 is modelled as *x*_i_(0) = *vY*_i_ for *i* = 1, …, *K*, where *Y*_i_ ∼ Po(*u*_i_/*v*) are independent Poisson random variables and *v* is a parameter modulating the level of the stochastic involved in the sampling.

Next, we aim to quantify the effects of inherent stochasticity in the agent-based model. Performing a finite-size expansion of the master equation associated with the model [35, 36], we derive a system of stochastic differential equations for the density of of cells and ***x*** = **X***/N*_0_, then for large but finite *N*_0_ we obtain the Langevin equation

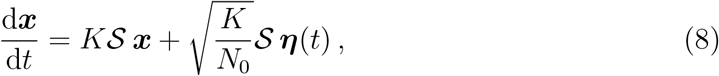

where **η**(*t*) is a *K*-dimensional white noise vector with correlator 𝔼[*η_i_*(*t*)*η_j_*(*t*^′^)] = *x_i_δ_ij_*δ(*t* − *t*^′^). The first term on the right describes the average behaviour of the model, and is the same as in Eq. 2. The second term captures the stochastic contributions arising from the finiteness of the population.

To gain more insight into the behaviour of this model, we start by writing down an Ornstein–Uhlenbeck (OU) model which approximates the behaviour of the Langevin equation as a stationary Gaussian process (see Section S.1 of Supplementary Materials). By employing standard results for stationary processes [36, 37] we obtain an expression for the time correlation matrix of **x** (see Section S.2 of Supplementary Materials). We then use the information gained from the OU process to calculate the standard deviation of *Q*(*t*), *σ*_Q_(*t*). Notice that by using the OU approximation, *Q*(*t*) is defined as a ratio of Gaussian distributed random variables which under certain conditions (discussed in Section S.3 of the Supplementary Materials) can be approximated by a Gaussian. Finally, we compute the envelope of two standard deviations about the mean of *Q*, defined as

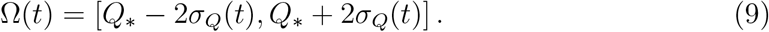

We denote with Ω the envelope obtained from the Langevin model with initial random sampling, with 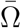 the envelope obtained from the system of Eq. 2 with initial random sampling and with Ω_**u**_ the envelope obtained from the of the Langevin model with the deterministic initial condition ***x***_0_ = ***u***. In the two panels of Fig. 4 we overlay 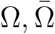 and Ω_***u***_, together with two numerical trajectories of *Q*: one (red) obtained by solving equation 8 and one (blue) by solving system of Eq. 2, both initialised with the same random initial condition.

**Fig. 4.**
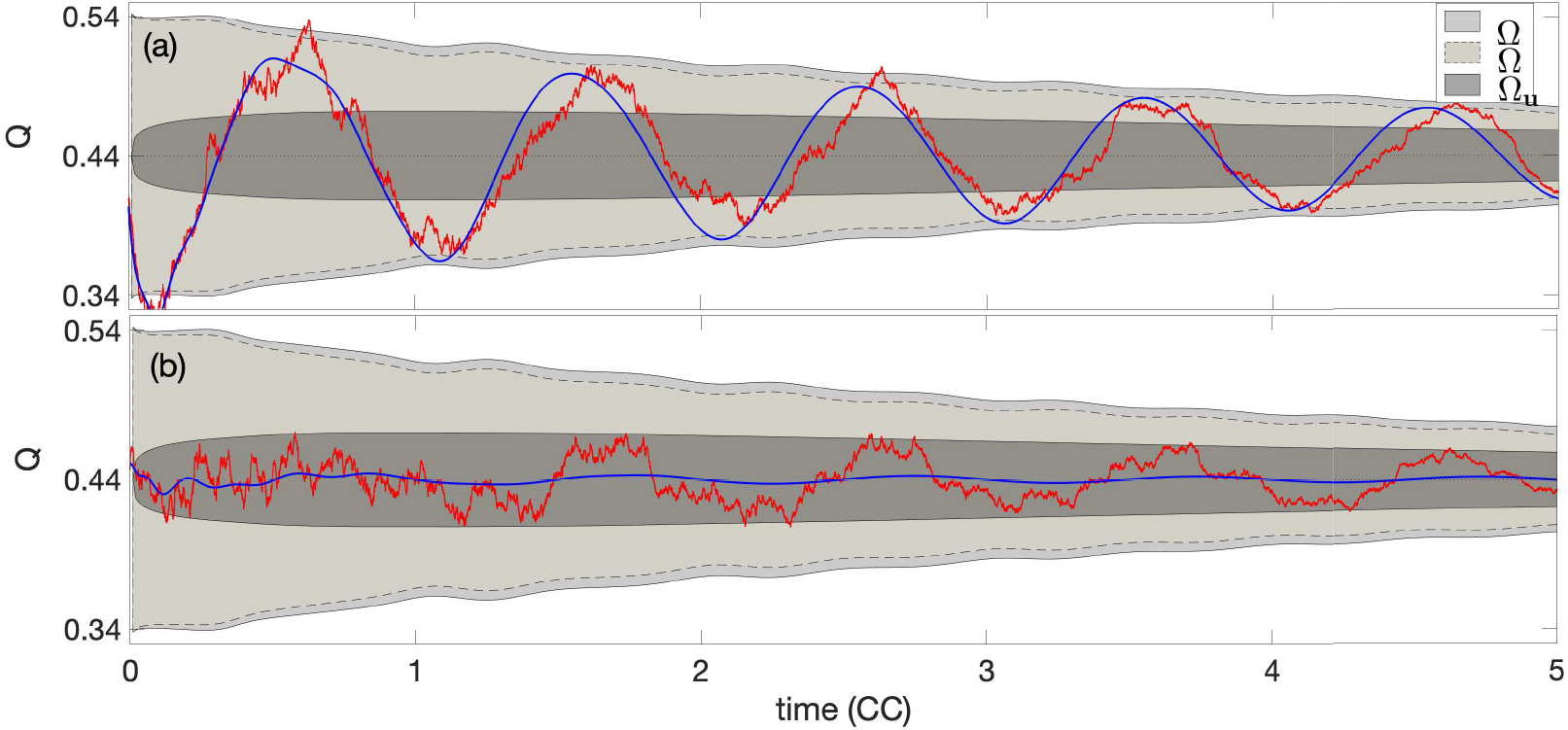
Finite-sample effects amplify the oscillations of *Q*. The two panels show the overlay of the three envelopes Ω (light grey region with solid line), 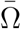 (medium grey region with dashed line) and Ω_***u***_ (dark grey region with solid line) together with two trajectories of *Q*(*t*) obtained by solving numerically (by using the Euler-Maruyama method with time step Δ*t* = 10^−3^) the Langevin model (red line) and the deterministic system of Eq. 2 (blue line) with the same, random initial condition. The two panels show two independent realisations of the models with different stochastic, initial conditions. The plots provide an example of two possible scenarios in which the intrinsic stochasticity of the Langevin model amplifies the oscillations of the deterministic system (panel a), or it triggers the emergence of new oscillations (panel b).

The results in Figure 4 show that in all three cases considered, accounting for the finite-sample stochasticity can lead to a persistent departure of *Q* from the equilibrium value. In the two cases which account for the initial random sampling, Ω and 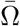, the envelopes present an evident initial departure from equilibrium, which is sustained for several cell-cycle times, halving after approximately four periods. The inherent dynamical stochasticity of the Langevin model tends to amplify the departure from the equilibrium as evident in Ω. Interestingly, both these envelopes have slightly fluctuating edges. In contrast, the envelope initialised at the invariant distribution, Ω_***u***_ shows an initial, fast expansion, followed by a phase of slower decay. Notice that Ω_***u***_ lies well inside Ω for all time. This suggests that the initial random sampling plays a role for the entire duration of the experiment. The numerical trajectories overlaid show good agreement with these findings. In particular, the solution of system of Eq. 2 (blue line) lies well inside 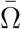 while the simulation of the Langevin equation (red line) shows a larger departure but it remains almost entirely inside the envelope Ω.

Notice that both trajectories in Fig. 4 (a) show clear oscillations about the origin with similar phases. The Langevin solution has increased oscillation amplitude in comparison to the solution of the deterministic system of Eq. 2. Although this amplification phenomenon, due to the stochasticity of the Langevin model, is common, we also find that, for some initialisations, the oscillations appear only in the Langevin model and not in the deterministic model as shown in Fig. 4 (b).

To quantify the appearance of the oscillations, we look at the time-autocorrelation function of *G*(*t*) which we define as

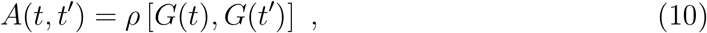

where *ρ* denotes the correlation coefficient (defined in Eq. (S.9) of Supplementary Materials) and can be computed using the formula for the correlation matrix (see Section S.2 of Supplementary Materials). Fig. 5 shows the evolution of the autocorrelation function, *A*(*t, t*^′^) as function of *t*^′^, for *t* = 0, 2 and 4, respectively. In each panel we plot *A*(*t, t*^′^) calculated analytically, using the correlation matrix, (black solid line) and the simulated value obtained by averaging 50 independent trajectories of the Langevin model (orange line). All three panels show a good agreement between the analytical formula and the simulated counterpart. Moreover, the results confirm the presence of strong fluctuations on the time autocorrelation of *G, i*.*e*. the number of cells in the G1-phase, with a period of exactly one cell cycle. As *Q* converges to the equilibrium,the amplitude of the oscillations decreases and the autocorrelation function *A*(*t, t*^′^) tends to unity.

**Fig. 5.**
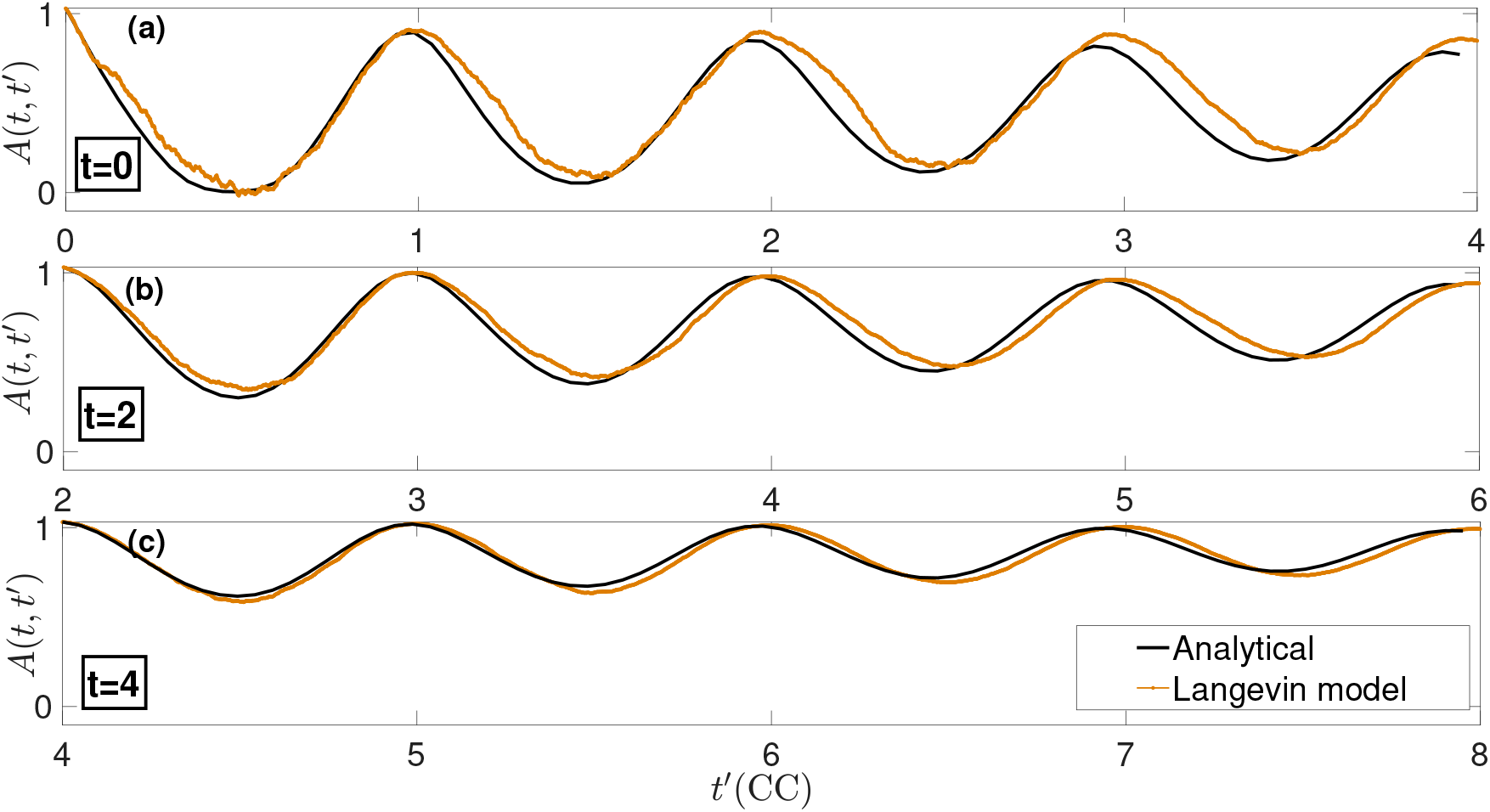
The time-autocorrelation function. The three panels (a), (b) and (c) show the time-autocorrelation function, *A*(*t, t*^′^) at time *t* = 0, 2 and 4, respectively, obtained analytically from the full stochastic model (back line) and by averaging over 50 independent simulations of the Langevin model (orange line). The parameters of the model are the same as in Fig. 1. Time is normalised with respect to the average cell-cycle time.

## IV. DISCUSSION

In this work we highlight the importance of demographic noise to the early dynamics of growing populations with non-exponentially distributed generation times. We demonstrate that finite-sample effects can recapitulate the synchronisation in the cell-cycle phase that we previously observed [16]. To provide insight in these observations we adopt a multi-stage approach to model both the total cell-cycle time distribution and the distribution of the G1 duration and we derive both a deterministic and a stochastic representation for the time evolution of the ratio *Q*. We find that the stochasticity in the initial sampling of cells can lead to a departure from the invariant distribution which triggers a transient oscillatory phase. The presence of intrinsic stochasticity in the dynamics tends to amplify these oscillations and delay their exponential decay.

We characterise the transient and asymptotic phases of the multi-stage model by deriving an analytical formula for the variance of the amplitude of the oscillations. By comparing our results with the experimental data from a proliferation assay of C8161 melanoma cells, we find that the frequency of the amplitude of the observed oscillations are consistent with those generated by demographic noise alone. Our findings suggest that finite-sample stochasticity plays a crucial role in the early stage dynamics of growing populations and that it can provide an explanation for observed synchronisation in the subpopulation of the cell cycle phases.

From a theoretical point of view, our study provides a further understanding of the relation between cell-cycle distribution and global population dynamics. Whilst our analysis employed a multi-stage model, the results of our analysis are amenable to extension to more general type of cell-cycle distributions. In fact, for certain choices of the model parameters, the Erlang distribution adopted in this paper is an excellent approximation of a Gaussian distribution. In Section S.5 of the Supplementary Materials we compute the relative entropy (Kullback-Leibler divergence) between Erlang and Gaussian distributions to show that the Erlang distribution tend to a Gaussian distribution as *K* → ∞. In principle, one could use this fact to study the applicability of our findings to Gaussian cell-cycle time and, hence, compare our results with other relevant studies which rely on a Gaussian approach [9, 10]. For example, Jafarpour [9] used a Gaussian model to study the connection between mother-daughter size regulation in bacteria and the decay of transient fluctuations. Our study focuses on synchrony emerging even in the absence of correlation of CCTs, we expect that accounting for such mechanisms will tend to amplify the amplitude of the oscillations predicted by the model, however, the analysis of this phenomenon is left for future study.

From an experimental point of view, our results highlight the routinely-overlooked importance of the sample size when performing experiments which involve small populations. In particular, any data interpretation should be carried out with the role played by finite-sample stochasticity in mind. Employing larger initial populations, for example by increasing the size of microscopy images, would diminish the amplitude of the synchronisation. However, since the oscillations described in this paper are an intrinsic phenomenon due to the finiteness of the population, the aim of any intervention would be to mitigate their effects, since they cannot be completely eradicated. Our analysis provides a novel platform to quantify the extent of finite-sample effects which can then be used to assess their relevance in experimental contexts.

One of the central implications of this study is the need for further experiments in order to understand and control the fluctuating phenomena observed in proliferating cell populations. A major aspect that remains unclear and that could serve as a guideline when designing new experimental protocols, is the spatial extent of the oscillatory phenomenon. Accounting for cell motility and spatial correlations of mother-daughter cells is likely to play an important role in the context of synchronising subpopulations [38]. In particular, we should expect cell motility to diminish spatial correlation, leading to faster decay of the oscillations of cells in a given window of space. In order to progress in this direction, we advocate that new experiments need to undertaken with increased microscopy field of view and larger cell populations. Similarly, the accompanying theoretical framework could be adapted by studying spatial extensions of the multi-stage models presented in this paper. Ultimately, a clearer understanding of the interplay of correlation of mother-daughter CCTs, cell motility and population size would help to identify the spatial scales at which the synchronisation phenomenon shows up and, in turn, those at which it is irrelevant.

While we primarily focus on explaining the synchronised oscillatory behaviour observed in subpopulations of melanoma cells in a proliferation assay, we stress that the general protocol developed here will be useful in order to determine the extent of effects due to finiteness of the cell population in a broad class of applications. Our proliferation assays are typical experimental protocols used to investigate the efficiency of cell-cycle-inhibiting drugs [4, 21], hence our findings may impact upon the reproducibility of such experiments, the efficacy of treatment protocols [22, 23] and the findings of mathematical models of these experiments [39–41]. Our work suggests that inherent synchronisation will also occur in bacterial populations and consequently that studies of bacterial pathogen growth may be impacted. Many experimental protocols rely on the synchronisation of cell populations in order to study the structural and molecular events that occur throughout the cell cycle, providing information about gene expression patterns, post transcriptional modification and contributing to drug discovery [24]. The improved understanding of the impacts of demographic noise on the evolution of synchronous populations showcased here will shed light on the potential impact that desynchronisation has on the results of these studies.

## AUTHOR CONTRIBUTIONS

E.G. designed the analytical tools, analysed the data and wrote the manuscript. S.T.V. performed the experiments. N.K.H. generated the FUCCI-C8161 cells. All authors provided feedback and gave approval for final publication.

## ACKNOWLEDGMENTS

E.G. was supported by the Santander Travel Award. N.K.H. is a Cameron fellow of the Melanoma and Skin Cancer ResearchInstitute, Australia, and is supported by the National Health and MedicalResearch Council (APP1084893). M.J.S. is supported by the Australian Research Council Discovery Program (DP170100474). TR acknowledges the support of the Royal Society. The authors declare no competing interests.

