## Supplementary Information for "Synchronised oscillations in growing cell populations are explained by demographic noise"

### Supplementary materials to accompany: "Synchronised oscillations in growing cell populations are explained by demographic noise"

Enrico Gavagnin [a],[1] Sean T. Vittadello,[2] Gency Gunasingh,[3] Nikolas K. Haass,[3] Matthew J. Simpson,[4] Tim Rogers,[5] and Christian A. Yates[5]

[1]*School of Biological Sciences*

*University of Bristol, Bristol, UK*

[2]*School of BioSciences*

*University of Melbourne, Melbourne, Victoria, Australia*

[3]*The University of Queensland, The University of Queensland Diamantina Institute, Brisbane, Queensland, Australia*

[4]*School of Mathematical Sciences*

*Queensland University of Technology, Brisbane, Queensland, Australia*

[5]*Department of Mathematical Sciences*

*University of Bath, Bath, UK*

This document contains the supplementary materials which accompany the paper Gavagnin et al. [1]. Sections S.1, S.2 and S.3 contain some details of the mathematical derivation of the analytical formula for the envelope of two standard deviation $Q$. In Section S.4 we explain the method adopted to parametrise our multi-stage model from the experimental images. Section S.5 contains the computation of the relative entropy between an Erlang and a Gaussian distribution. Section S.6 contains supplementary figures. All accompanying codes of the publication (MATLAB 2019b) can be found at https://github.com/EnricoGavagnin/sync_multi_stage.git

#### S.1. THE ORNSTEIN-UHLENBECK APPROXIMATION

We can simplify the Langevin model given by Eq. 8 of the main document by replacing the dependence on $\boldsymbol{x}$ in the correlator of $\boldsymbol{\eta}(t)$ with $\mathbb{E}[\boldsymbol{x}] = K\boldsymbol{u}e^{K\lambda t}$. The resulting equation consists of the high-dimensional non-autonomous Ornstein-Uhlenbeck (OU) process

$$\frac{\mathrm{d}\hat{\boldsymbol{x}}}{\mathrm{d}t} = K\mathcal{S}\hat{\boldsymbol{x}} + K\sqrt{\frac{e^{K\lambda t}}{N_0}}\mathcal{S}\boldsymbol{\psi}(t)\,, \tag{S.1}$$

where $\boldsymbol{\psi}(t)$ is a $K$-dimensional white noise vector with correlator $\mathbb{E}[\eta_i(t)\eta_j(t')] = u_i\delta_{ij}\delta(t-t')$. We explore the behaviour of the two models, the OU process given by Eq. S.1, and the Langevin equation 8 of the main document in Fig. S.1. The results suggest that the OU process is an accurate approximation of the Langevin equation, in particular the presence of the oscillations is evident in both the modelling regimes (Fig. S.1(a)). In Fig. S.1(b), we compare the distributions of $Q(t)$ at $t = 1, 3$ and $5$ obtained by averaging over 1000 independent simulations, which show good agreement between the two models.

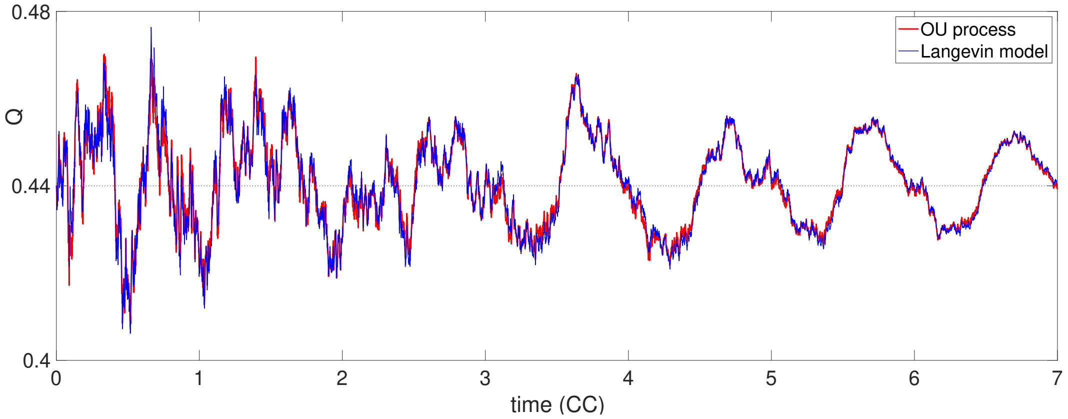


(a)

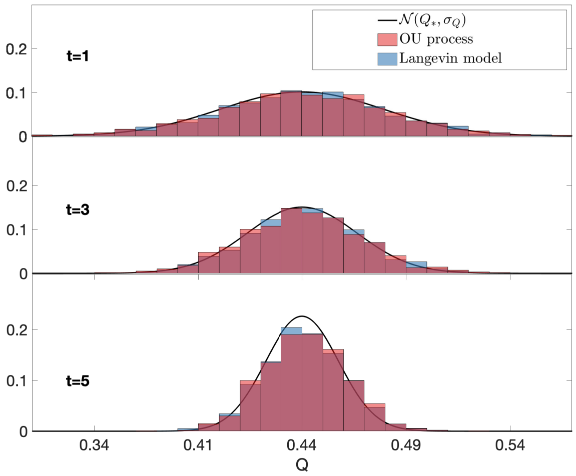


(b)

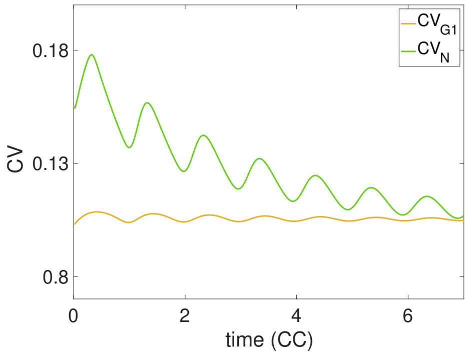


(c)

Fig. S.1. Comparison of OU process and the Langevin equation model. Panel (a) shows two evolutions of $Q$ for the OU process (red) and the Langevin equation (blue). The two trajectories are realised using Euler-Maruyama method with time step $\Delta t = 10^{-3}$ and the same randomly generated numbers. In panel (b) we plot the distribution of $Q$ at three time points ($t = 1$, $t = 3$ and $t = 5$). The two overlaid histograms represent the distributions of 1000 independent simulations of the OU process (red) and the Langevin Equation (blue). The black line represent the distribution $N(Q_*, \sigma_Q(t))$. All the parameters are the same as Fig. 2 of the paper. Panel (c) shows the values of $CV_{G1}(t)$ (green line) and $CV_N(t)$ (yellow line) indicating that $Q$ will be Gaussian distributed for the chosen parameter values. The parameters are the same as Fig. 2 of the paper and time is normalised with reference to the average cell-cycle time.

#### S.2. THE CORRELATION MATRIX

For a stochastic initial condition, $\boldsymbol{x}_0$, as described in Section III C of the main document, we can compute the correlation matrix at time $t = 0$, as

$$C_0 = \mathbb{E}\left[x_i(0)x_j(0)\right] = \begin{cases} u_i u_j & \text{for } i \neq j \\ u_i^2 + \frac{u_i}{v} & \text{for } i = j \end{cases} . \tag{S.2}$$

We can rewrite this as $C_0 = \boldsymbol{u}\boldsymbol{u}^T + \frac{1}{v}M$, where $M = Diag(\boldsymbol{u})$.

We then focus on computing the correlation matrix $C(t,t') = \mathbb{E}\left[\hat{\boldsymbol{x}}(t)\,\hat{\boldsymbol{x}}^T(t')\right]$ for the OU process S.1, as an approximation for the correlation matrix of the Langevin model. By applying general results for OU processes (See Section 4.5 of [2]) we have:

$$C(t,t') = e^{Kt\mathcal{S}} C_0 e^{Kt'\mathcal{S}^T} + \frac{K}{N_0}\int_0^{\min(t,t')} e^{K(t-\tau)\mathcal{S}}\mathcal{S}\left(Me^{K\lambda\tau}\right)\mathcal{S}^T e^{K(t'-\tau)\mathcal{S}^T}\,d\tau\,. \tag{S.3}$$

We can use expression 5 of the main document and the fact that $e^{Kt\mathcal{S}}\boldsymbol{u}\boldsymbol{u}^T e^{Kt'\mathcal{S}^T} = \boldsymbol{u}\boldsymbol{u}^T e^{K(t+t')\lambda}$, to write down the $(i,j)$ element of S.3 for $t < t'$ as

$$\begin{aligned} C_{i,j}(t,t') =& u_i u_j e^{K(t+t')\lambda} + \frac{1}{vK^2}\sum_{k,l,m=1}^{K}\frac{1}{(1+\lambda_k)^{i-m}}\frac{1}{(1+\lambda_l)^{j-m}}\frac{2\lambda}{(1+\lambda)^m}e^{K(t\lambda_k+t'\lambda_l)} \\ &+ \frac{1}{N_0K}\sum_{k,l,m=1}^{K}\frac{\lambda_k}{(1+\lambda_k)^{i-m}}\frac{\lambda_l}{(1+\lambda_l)^{j-m}}\frac{2\lambda}{(1+\lambda)^m}\int_0^t e^{K[(t-\tau)\lambda_k+(t'-\tau)\lambda_l+\lambda\tau]}\,. \end{aligned} \tag{S.4}$$

Substituting the expressions 3 of the main text for $\boldsymbol{u}^k$ and $\boldsymbol{v}^k$, using the formula $\sum_{m=1}^{K}(1+\lambda_k)^m(1+\lambda_l)^m/(1+\lambda)^m = (1+\lambda_k)(1+\lambda_l)/[(1+\lambda_k)(1+\lambda_l)-(1+\lambda)]$ and upon rearranging terms, we obtain

$$
\begin{aligned}
C_{i,j}(t,t') =& \frac{4\lambda^2}{(1+\lambda)^{i+j}} e^{K(t+t')\lambda} \\
&+ \frac{2\lambda}{K^2} \sum_{k,l=1}^{K} \frac{(1+\lambda_l)^{1-j}(1+\lambda_k)^{1-i}}{(1+\lambda_k)(1+\lambda_l)-(1+\lambda)} \Bigg[ \frac{1}{v} e^{K(t\lambda_k + t'\lambda_l)} \\
&- \frac{\lambda_k \lambda_l}{N_0(\lambda - \lambda_k - \lambda_l)} \left( e^{K(t\lambda_k + t'\lambda_l)} - e^{K((t'-t)\lambda_l + t\lambda)} \right) \Bigg] .
\end{aligned}
\tag{S.5}
$$

It should be noted that alternative techniques could be employed to approximate the correlation matrix of the Langevin model. For instance, it is possible to write down the equation for the second and third moment of $\boldsymbol{x}$ directly from the Langevin equation (Eq. 8 of the main document) and to employ a moment closure approximation in order to truncate the dependency on the fourth moment. For reasonable choices of closure approximations, we expect such an approach would lead to similar formulae to those we obtained in this section by using the OU approximation.

#### S.3. THE ENVELOPE OF TWO STANDARD DEVIATIONS OF $Q$

We recall the definition of the envelope of two standard deviations of $Q(t)$ as $\Omega(t) = [Q_* - 2\sigma_Q(t), Q_* + 2\sigma_Q(t)]$, where $\sigma_Q(t)$ denotes the standard deviation of $Q(t)$. To compute $\sigma_Q$ we employ the OU approximation (see Section S.1). From a fixed initial condition, the solutions of S.1 evolve as a Gaussian process with mean $\bar{\boldsymbol{x}}(t)/N_0$. We can write $G(t) \sim \mathcal{N}(\mu_G(t), \sigma_G(t))$ and $N(t) \sim \mathcal{N}(\mu_N(t), \sigma_N(t))$ where

$$
\mu_G(t) = Q_* e^{K\lambda t}, \qquad \sigma_G^2(t) = \sum_{i,j=1}^{\alpha K} C_{i,j}(t,t) - Q_*^2 e^{2K\lambda t}, \tag{S.6a}
$$

$$
\mu_N(t) = e^{K\lambda t}, \qquad \sigma_N^2(t) = \sum_{i,j=1}^{K} C_{i,j}(t,t) - e^{2K\lambda t}. \tag{S.6b}
$$

Notice that, $Q$ is defined as a ratio between two Gaussian distribution and, in general, this does not imply that $Q(t)$ is Gaussian. However, Hayya et al. [3] showed that the ratio of two Gaussian can be well approximated as a Gaussian, under certain conditions on the *coefficient of variation* (CV) of the numerator and denominator. Precisely, provided that

$$CV_N = \frac{\sigma_N}{\mu_N} < 0.39 \qquad \text{and} \qquad CV_{G1} = \frac{\sigma_{G1}}{\mu_{G1}} > 0.005 \tag{S.7}$$

Hayya et al. [3] demonstrate that $Q$ is close to a Gaussian distribution. Moreover, we can approximate the variance of $Q$ by Taylor expanding to the second order which leads to

$$\begin{aligned} \sigma_Q^2 &\approx \sigma_N^2 \frac{\mu_G^2}{\mu_N^4} + \frac{\sigma_G^2}{\mu_N^2} - 2\rho\mu_G \frac{\sigma_N \sigma_G}{\mu_N^3} \\ &= \frac{1}{\mu_N^2}\left[\sigma_N^2 Q_*^2 + \sigma_G^2 - 2\sigma_N\sigma_G Q_*\,\rho\left[G,N\right]\right], \end{aligned} \tag{S.8}$$

where $\rho$ denotes the correlation coefficient, defined as

$$\rho\left[Y_1, Y_2\right] = \frac{\mathbb{E}\left[Y_1 Y_2\right] - \mathbb{E}\left[Y_1\right]\mathbb{E}\left[Y_2\right]}{\sqrt{\text{Var}\left[Y_1\right]\ \text{Var}\left[Y_2\right]}}\,. \tag{S.9}$$

Notice that we can compute $\mathbb{E}\left[G(t)N(t)\right]$ in Eq. S.9 in terms of the correlation matrix $C$ as

$$\mathbb{E}\left[G(t)N(t)\right] = \sum_{i=1}^{K}\sum_{j=1}^{\alpha K} C_{i,j}(t,t)\,.$$

Finally we check that the conditions S.7 are satisfied for our parameter choices. In Fig. S.1 (c) we report the value of $CV_{G1}$ (yellow line) and $CV_N$ (green line) for the parameter values obtained in Section S.4. The two coefficients lie well inside the range indicated by Hayya et al. [3], suggesting that the ratio $Q$ is close to a Gaussian distribution for the parameters considered. Notice that the plots in Fig. S.1(b) provide further confirmation of this by showing good agreement between the distribution of $Q$ and the Gaussian distribution $\mathcal{N}(Q_*, \sigma_Q(t))$.

#### S.4. PARAMETER INFERENCE

To infer the parameters of the multi-stage model, we simultaneously fit the distribution of the total cell-cycle time and of the G1 duration of 200 randomly selected cells.

Let $\boldsymbol{H}_T$ and $\boldsymbol{H}_{G1}$ denote the histogram representations of the probability density function (pdf) of the total cell-cycle time and the G1 duration, respectively, with a bin width of one hour. For example, $(\boldsymbol{H}_T)_i$ denotes the proportion of cells with a cell-cycle time in the interval $[ih, (i+1)h)$. We denote with $\boldsymbol{H}_{E(K,\beta)}$ the histogram obtained by discretising an Erlang distribution with parameters $(K, \beta)$ with the same bin width, *i.e.* $(\boldsymbol{H}_{E(K,\beta)})_i = \frac{\beta^K}{(K-1)!}\int_i^{i+1} x^{K-1}e^{-\beta x}dx$.

For a given combination of parameters, $(K, \beta, \alpha)$, one can consider the distance measure

$$I(K, \beta, \alpha) = \|\boldsymbol{H}_T - \boldsymbol{H}_{E(K,\beta)}\|_1 + \|\boldsymbol{H}_{G1} - \boldsymbol{H}_{E(\alpha K,\beta)}\|_1 \,, \qquad \text{(S.10)}$$

where $\| - \|_1$ denotes the 1-norm. To determine the parameter combination which provides the best simultaneous fit of the two distribution, we evaluated the function $I$ in the parameter range $K \in [10, 150]$ $\beta \in [1, 10]$ and $\alpha \in [0, 1]$. We find that the combination $K^* = 92$, $\beta^* = 4.96$ and $\alpha^* = 33/92$ minimises the statistic $I$ in the parameter region considered and, hence, we select these parameters for the multi-stage model.

In order to determine the level of initial stochasticity of the model, we choose the value of the parameter $v$ which provides the best fit between the experimental standard deviation and the one predicted from the model (see Figure 2 (b) of the main text).

This is done by minimising the 1-norm of the distance between the experimental time series of $\sigma_Q(t)$, obtained from the 30 experimental trajectories, and the corresponding theoretical time series obtained using formula S.5. After performing a direct parameter exploration for $v \in \{\delta_v k \,|\delta_v = 0.1,\ k = 1 \ldots 10^3\}$ we found that the distance between the two trajectories is minimised by choosing $v = 94.3$, hence we select this value as parameter for the model. The initial average population size, $N_0$ is chosen to match the experimental value, averaged over the 30 repeats, $N_0 \approx 381$.

#### S.5. THE KULLBACK LEIBLER DIVERGENCE BETWEEN ERLANG AND GAUSSIAN DISTRIBUTION

We compute the relative entropy (Kullback-Leibler divergence, $D_{KL}$) between an Erlang and a Gaussian distribution as a measure of the distance between the two distributions.

For two distributions, $p(x)$ and $q(x)$, the KL divergence is defined as:

$$D(p,q) = \int_{-\infty}^{\infty} p(x) \log\left[\frac{p(x)}{q(x)}\right] dx\,. \tag{S.11}$$

We set $p(x)$ to be the probability density function (pdf) of an $Erlang(K,\beta)$ and $q(x)$ to be the pdf of a Gaussian with same mean and variance, *i.e.* $\mathcal{N}\left(\frac{K}{\beta}, \frac{K}{\beta^2}\right)$. We then obtain

$$
\begin{aligned}
D(K,\beta) =& D\left(p\left(K,\beta\right), q\left(\frac{K}{\beta},\frac{K}{\beta^2}\right)\right) \\
=& \frac{\beta^K}{(K-1)!}\int_0^\infty x^{K-1}e^{-\beta x}\log\left[\frac{\beta^{K-1}\sqrt{2\pi K}}{(K-1)!}x^{K-1}e^{\frac{\beta^2}{2K}\left(x-\frac{K}{\beta}\right)^2-\beta x}\right]dx \\
=& \log\left[\frac{\beta^{K-1}\sqrt{2\pi K}}{(K-1)!}\right]\frac{\beta^K}{(K-1)!}\int_0^\infty x^{K-1}e^{-\beta x}dx \\
&+ (K-1)\frac{\beta^K}{(K-1)!}\int_0^\infty x^{K-1}\log(x)e^{-\beta x}dx \\
&+ \frac{\beta^K}{(K-1)!}\int_0^\infty\left[-\beta x+\frac{\beta^2}{2K}\left(x-\frac{K}{\beta}\right)^2\right]x^{K-1}e^{-\beta x}dx\,.
\end{aligned}
\tag{S.12}
$$

Notice that the first integral of Eq. S.12 is exactly the pdf of an Erlang distribution, which simplifies to unity. The second and third integral in S.12 require more work. By using integration by parts and upon simplification, we get to the final expression

$$
D(K,\beta) = \log\left[\frac{\beta^{K-1}\sqrt{2\pi K}}{(K-1)!}\right] + (K-1)\left(\mathcal{H}_{K-1}-\log(\beta)-\gamma\right) - K + \frac{1}{2}\,, \tag{S.13}
$$

where $\mathcal{H}_{K-1} = \sum_{i=1}^{K-1}\frac{1}{i}$ is the $(K-1)$-th harmonic number and $\gamma = \lim_{n\to+\infty}\left(\sum_{i=1}^{n}\frac{1}{i}-\log(n)\right)$ denotes the Euler-Mascheroni constant.

Using the expression of Eq. S.13 it is possible to show that $D(K,\beta)$ is a decreasing function of $K$ and $D(K,\beta)\sim\mathcal{O}(K^{-1})$ for $K\to+\infty$. This is not a surprise, since by central limit theorem we know that the Erlang distribution converges to a Gaussian with same mean and variance. Since the CV of an Erlang$(K,\beta)$ is given by $K^{-\frac{1}{2}}$, we can rephrase by saying that the KL divergence scales proportionally to the square of the CV of the Erlang distribution.

Fig. S.2 shows the plot of $D(K,\beta)$ with $\beta = K$ for different values of $K$. In the overlaid panels the two distributions are compared for $K = 5, 10, 20, 30$ and 60. The results

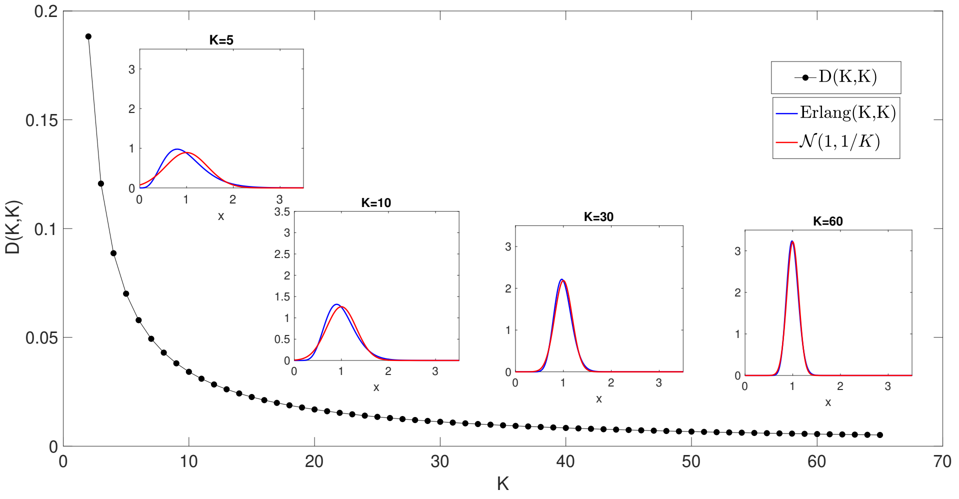


Fig. S.2. The Kullback Leibler (KL) divergence between an Erlang distribution and Gaussian distribution. The black dotted line in the main panel shows the KL divergence between an Erlang distribution of parameters $(K, K)$ and a Gaussian distribution of parameters $(1, 1/K)$ as function of $K$. The four overlaid panels show the comparison of the two distributions, Erlang (blue) and Gaussian (Red), for $K = 5, 10, 30$ and 60 (from left to right).

highlight the good level of similarity between the Erlang and Gaussian distributions for large $K$ - small values of the CV. For example, for $K > 25$, *i.e.* CV$< 0.2$, we have $D(K, K) < 0.02$ which corresponds to good agreement between the two distributions.

#### S.6. SUPPLEMENTARY FIGURES

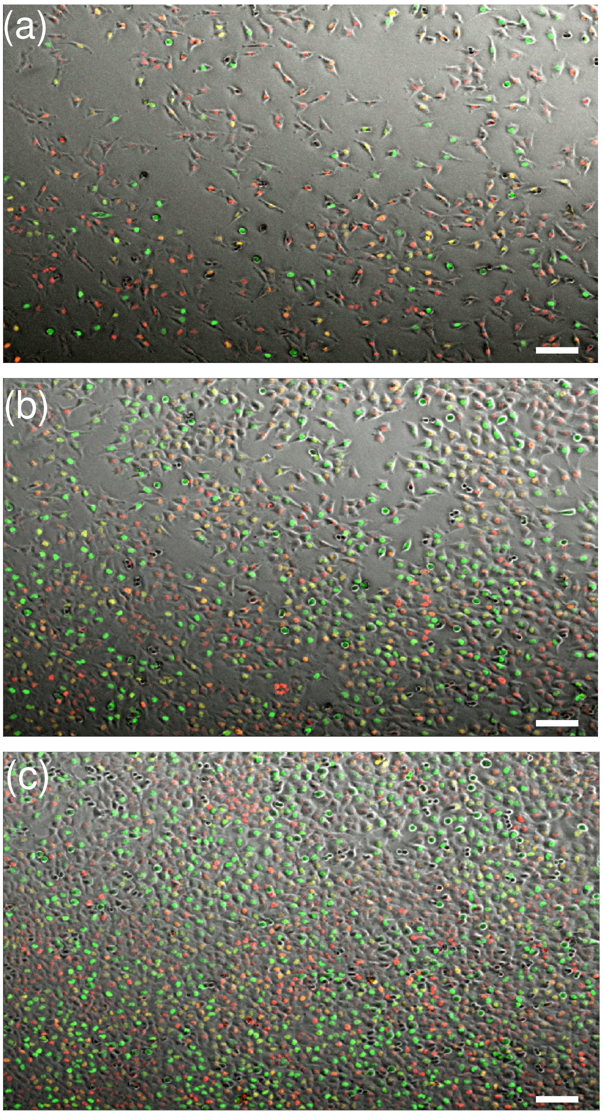


Fig. S.3. Microscopy images of proliferation assay. Panels (a) to (c) show three snapshots of an epifluorescence microscopy image time series - merged images of bright-field and the red and green fluorescence channels. Panel (a) is taken at the beginning of the recording, $t = 0$ h, panel (b) is taken half-way through the recording, $t = 24$ h, and at panel (c) at the end of the recording, $t = 48$ h (scale bar $100 \mu m$). The cells with red nuclei are in the G1 (gap 1) phase of the cell cycle, the ones with yellow nuclei are in the eS (early-synthesis) phase and those with green nuclei are in one of the remaining consecutive phases: S (synthesis), G2 (gap 2) or M (mitosis).

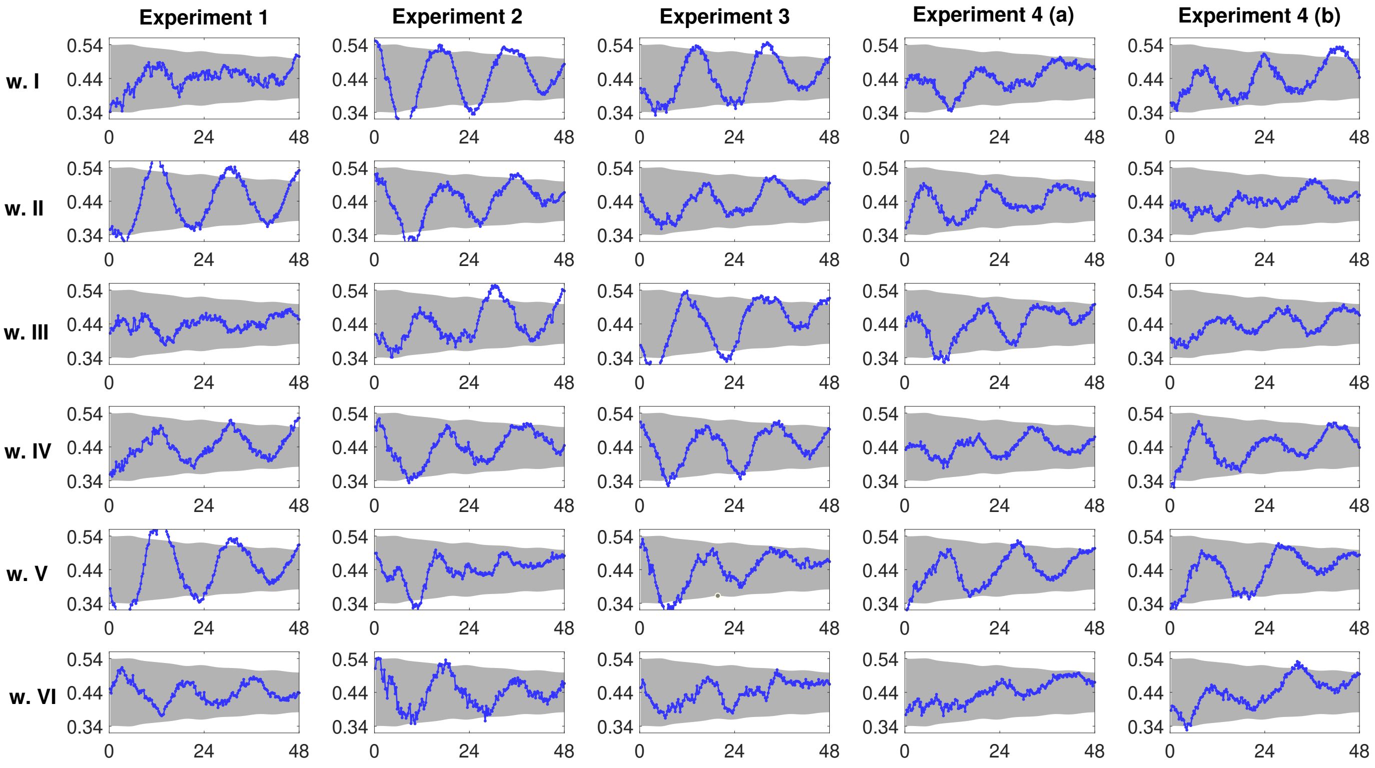


Fig. S.4. Comparison 30 time series obtained from the data (blue lines), together with the envelope of two standard deviations, $\Omega$ (light grey regions) predicted using the multi-stage model. The parameters of the multi-stage models are obtain by fitting the distribution of the total cell-cycle time and G1 duration (see Section S.4): $K = 92, \alpha K = 33, \beta = 4.96\text{h}^{-1}, v = 94.3$ and $N_0 = 381$.

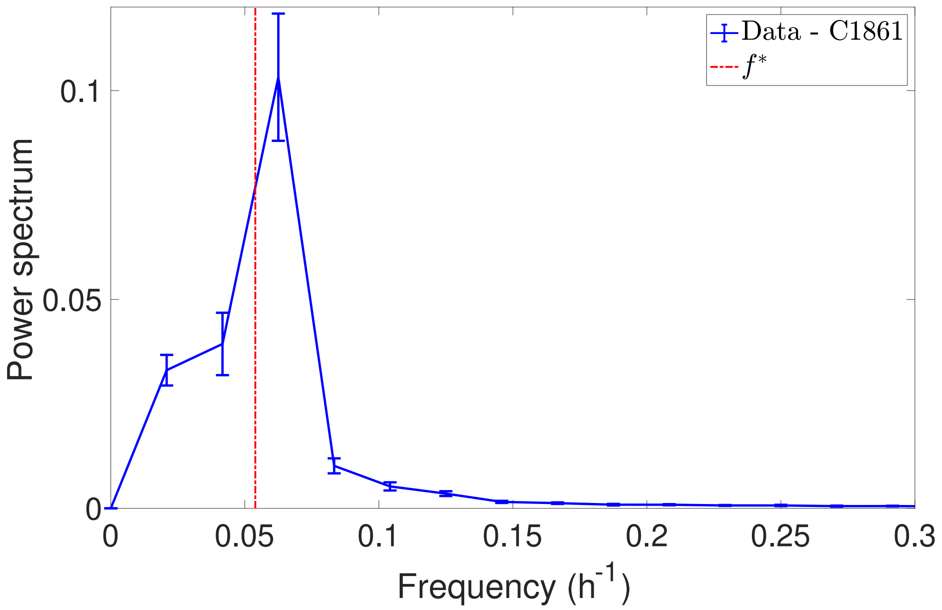


Fig. S.5. Average Fourier power spectrum analysis of the time series of $Q(t)$. The plot shows the experimental Fourier power spectrum of $Q(t)$ averaged over the 30 time series (blue lines - bars denote interval of one standard deviation). The red line highlights the dominant frequency predicted from the multi-stage model, corresponding to one CCT, $f^* = \beta/K = 0.054\,\mathrm{h}^{-1} = (18.5\ \mathrm{h})^{-1}$.

[1] E. Gavagnin, S.T Vittadello, G. Gunasingh, N.K Haass, M.J. Simpson, T. Rogers, and C.A. Yates. Synchronised oscillations in growing cell populations are explained by demographic noise. *t.b.a.*, 2020.

[2] C. Gardiner. *Stochastic methods*, volume 4. Springer Berlin, 2009.

[3] J. Hayya, D. Armstrong, and N. Gressis. A note on the ratio of two normally distributed variables. *Manag. Sci.*, 21(11):1338–1341, 1975.
